# Comprehensive investigation and regulatory function of lncRNAs engaged in western honey bee larval immune response to *Ascosphaera apis* invasion

**DOI:** 10.1101/2022.10.28.514173

**Authors:** Yaping Ye, Xiaoxue Fan, Qi Long, Jie Wang, Wende Zhang, Zongbing Cai, Minghui Sun, Xiaoyu Gu, Peiyuan Zou, Dafu Chen, Rui Guo

## Abstract

*Ascosphaera apis* is a fungal pathogen that exclusively infects bee larvae, causing chalkbrood disease, which results in severe damage for beekeeping industry. Long non-coding RNAs (lncRNAs) are versatile regulators in various biological processes such as immune defense and host-pathogen interaction. However, expression pattern and regulatory role of lncRNAs involved in immune response of bee host to *A. apis* invasion is still very limited. Here, *Apis mellifera ligustica* 4-, 5-, and 6-day-old larvae inoculated by *A. apis* spores (AmT1, AmT2, and AmT3 groups) and corresponding un-inoculated larval guts (AmCK1, AmCK2, and AmCK3 groups) were prepared and subjected to deep sequencing, followed by identification of lncRNAs, analysis of differentially expressed lncRNAs (DElncRNAs), and investigation of DElncRNA-regulated host immune responses. In total, 3 746 *A. m. ligustica* lncRNAs were identified, including 78 sense lncRNAs, 891 antisense lncRNAs, 1 893 intergenic lncRNAs, 346 bidirectional lncRNAs, and 210 intronic lncRNAs. Compared with corresponding un-inoculated larval guts, 357, 236, and 505 DElncRNAs were discovered in the *A. apis*-inoculated 4-, 5-, and 6-day-old larval guts. Additionally, there were 32 shared DElncRNAs. In AmCK1 vs AmT1, AmCK2 vs AmT2, and AmCK3 vs AmT3 comparison groups, 217, 129, and 272 DElncRNAs were respectively predicted to regulate neighboring genes via *cis*-acting manner, and these targets were associated with a series of GO terms and KEGG pathways of great importance, such as response to stimulus and Jak-STAT signal pathway. Moreover, competing endogenous RNA (ceRNA) network analysis indicated that 197, 95, and 356 DElncRNAs in the aforementioned three comparison groups could target 10, eight, and 21 DEmiRNAs and further target 147, 79, and 315 DEmRNAs, forming complex regulatory networks. Further investigation suggested that these targets were relevant to several key cellular and humoral immune pathways, such as phagosome and MAPK signaling pathway. Ultimately, the expression trends of nine randomly selected DElncRNAs were verified by RT-qPCR, indicative of the authenticity and reliability of our transcriptome data. Findings in this current work not only provide candidate DElncRNAs for functional study, but also lay a foundation for unclosing the mechanism underlying DElncRNA-regulated larval immune responses to *A. apis* infection.

## Introduction

Western honey bees (*Apis mellifera*) are widely reared in apiaries worldwide and produce a variety of honey bee products, such as honey, royal jelly, and propolis. *A. mellifera* pollinates a substantial quantity of crops and wild flowers, playing a critical role in ecological balance and food security (Potts et al., 2010). *Ascosphaera apis* is a widespread fungal pathogen that infects bee larvae and causes chalkbrood disease, leading to a sharp reduction in colony population and productivity (Aronstein and Murray, 2010). In the last two decades, researchers have conducted an array of studies on the interaction between honey bee larvae and *A. apis* at the morphological, ethological, or biochemical level (Guo et al., 2019; Chaimanee et al., 2017; Garrido-Bailón et al., 2013). Our group previously systematically investigated the *Apis mellifera ligustica* larval response and *A. apis* infection during chalkbrood (Chen et al., 2017a). For example, Chen et al. (2017a) revealed the cellular and humoral immune responses of western honey bee larvae to *A. apis* invasion based on deep sequencing combined with bioinformatics; Chen et al. (2017b) deciphered the transcriptomic dynamics of *A. apis* during the infection process in *A. m. ligustica* larvae.

Long noncoding RNAs (lncRNAs) are ncRNAs with a length of more than 200 nt and two or more exons (Zhang et al., 2019). Similar to mRNA structures, most lncRNAs are transcribed by RNA polymerase II and hence contain the 5’ cap and the 3’ polyA tail (Li et al., 2012). According to the location with respect to protein-coding genes, lncRNAs can be classified into sense lncRNAs, antisense lncRNAs, intergenic lncRNAs, intron lncRNAs, and other lncRNAs (Jarroux et al., 2017). LncRNAs are capable of exerting functions in versatile manners, namely, *cis*-acting regulation, *trans*-acting modulation, miRNA precursor, competing endogenous RNA (ceRNA) network, and translation into protein (Gil and Ulitsky, 2020). Increasing evidence suggests that lncRNAs are pivotal regulators in a large number of biological processes, including dose compensation (Carmona et al., 2018), cell cycle (Ma et al., 2021), and host-pathogen interaction (Carnero et al., 2016).

With the rapid development of next-generation sequencing technologies, abundant lncRNAs have been identified in animals, plants, and microorganisms (Fan et al., 2021; Zhou et al., 2021a; Guo et al., 2018a). In insects, lncRNA-associated studies have mainly focused on model insects such as fruit flies (*Drosophila*) and silkworms (*Bombyx mori*); however, knowledge of other insects, including honey bees, is still limited at present. Zhou et al. (2021b) discovered that *lncRNA-CR11538* in *Drosophila* restored Toll immunity homeostasis via interaction with the transcription factor Dif/Dorsal, further participating in host innate immunity. Previous work demonstrated that lncRNAs in honey bees were potentially involved in behavior (Feng et al., 2022), caste development (Humann et al., 2013), and host-pathogen/parasite interaction (Chen et al., 2019a). Our team previously deciphered the responses of both *A. m. ligustica* and *Apis cerana* larvae to *A. apis* invasion at the mRNA and miRNA levels (Chen et al., 2017a; Guo et al., 2019a). Accumulating evidence has demonstrated that ncRNAs (mRNAs, lncRNAs, circRNAs, etc.) containing miRNA response elements (MREs) could target mRNAs via competition with miRNAs, further regulating downstream gene expression and biological processes (Ala, 2020). However, no research has reported on the expression profile and regulatory role of lncRNAs during *A. m. ligustica* bee larval growth in response to *A. apis* invasion.

In the current work, 4-, 5-, and 6-day-old *A. m. ligustica* larval guts inoculated with *A. apis* spores and un-inoculated larval guts were subjected to strand-specific cDNA library construction and deep sequencing, followed by identification of lncRNAs and investigation of the expression profile. The potential regulatory functions of host DElncRNAs were then analyzed in combination with previously obtained small RNA-seq (sRNA-seq) data. To our knowledge, this is the first report of a lncRNA-regulated response of honey bee larvae to fungal invasion. Our findings will shed light on the mechanism underlying the lncRNA-mediated larval response to *A. apis* infection and offer novel insights into the interaction between *A. m. ligustica* larvae and *A. apis*.

## Materials and method

### Bee and fungus

*A. m. ligustica* colonies were reared in the teaching apiary of the College of Animal Sciences (College of Bee Science) at Fujian Agriculture and Forestry University. *A. apis* spores were prepared following the method developed by Jensen et al. (2013) and stored in the Honey Bee Protection Lab, College of Animal Sciences (College of Bee Science), Fujian Agriculture and Forestry University.

### Experimental inoculation and sample preparation

In our previous study, *A. m. ligustica* larvae were reared in 48-well culture plates in a constant temperature and humidity incubator (Shanghai Yiheng Scientific Instrument Co., Ltd.) as described by Peng et al. (1992). Briefly, the diet was mixed and frozen in smaller aliquots and preheated to 34°C before feeding; 2-day-old larvae were removed from the combs with a transferring tool to 10 μL diet; 3-day-old larvae (n=9) in the treatment group were fed 20 μL diet containing *A. apis* spores with a final concentration of 10^7^ spores/mL and then fed once a day with 30 μL (4-day-old), 40 μL (5-day-old), and 50 μL (6-day-old) diet; 3-day-old larvae (n=9) in the control group were fed once a day with 20 μL (3-day-old), 30 μL (4-day-old), 40 μL (5-day-old), and 50 μL (6-day-old) diet without *A. apis* spores; gut tissues of 4-, 5-, and 6-day-old larvae were harvested utilizing our previously developed protocol (Guo et al., 2019b) and then frozen in liquid nitrogen and kept at -80°C until deep sequencing and molecular experiments. Gut samples of 4-, 5-, and 6-day-old larvae in the control groups were named the AmCK1 group, AmCK2 group, and AmCK3 group, while those in the treatment groups were named the AmT1 group, AmT2 group, and AmT3 group, respectively.

### RNA extraction, cDNA library construction, and deep sequencing

The total RNA of the gut samples in the abovementioned six groups was extracted using the TRIzol method (Promega, USA). Oligo (dTs) were used to isolate poly (A) mRNA, which was subsequently fragmented and reverse transcribed using random primers (QIAGEN, Germany). Next, second-strand cDNA was synthesized with RNase H and DNA polymerase I, and the double-strand cDNA was then purified by the QiaQuick PCR extraction kit (QIAGEN, Germany). After agarose gel electrophoresis, the required fragments were purified using a DNA extraction kit (QIAGEN, Germany) followed by enrichment through PCR amplification (NEB, USA). The reaction conditions were set as follows: 98°C for 30 s, followed by 13 cycles of 98°C for 10 s and 65°C for 75 s, and 65°C for 5 s. Ultimately, the 6 cDNA libraries were subjected to deep sequencing by Guangzhou Gene Denovo Biotechnology Co., Ltd. using the Illumina HiSeqTM 4000 platform. Raw data generated from strand-specific library-based RNA-seq were deposited in the NCBI Sequence Read Archive (SRA) database and linked to BioProject number PRJNA406998.

### Quality control of raw data

Fastp software (version 0.18.0) (Chen et al., 2018) was used to perform quality control on the raw data by removing reads that contained adapters, more than 10% N-base or low-quality reads (Q-value≤20) to obtain high-quality clean reads. The obtained clean reads were mapped to the reference genome of *A. mellifera* (Amel_HAV3.1) using HISAT2 software (Kim et al., 2015) with default parameters.

### Identification of lncRNAs

The transcripts were assembled using a combination of Cufflinks (Trapnell et al., 2012) and TopHat2 software (Kim et al., 2013) and then aligned to the *A. mellifera* reference genome (Amel_HAV3.1) with Cuffcompare software (Meana et al., 2010) to detect novel transcripts. Transcripts with one of the classcodes “u, i, j, x, c, e, o” were defined as novel transcripts. Next, lncRNAs were filtered out according to the following criteria: (1) length ≥ 200 bp and (2) exon number ≥ 2. Furthermore, both CPC (Kong et al., 2017) and CNCI software (Sun et al., 2013) were used to predict the coding ability of the above lncRNAs, and only those without coding ability were regarded as reliable lncRNAs. The numbers of various types of lncRNAs, including intergenic lncRNAs, sense lncRNAs, and antisense lncRNAs, were calculated.

### Analysis of DElncRNAs

Following the method described by Chen et al. (2017a), the transcript expression level was normalized with the FPKM (fragments per kilobase of transcript per million mapped reads) method, which can eliminate the influence of different transcript lengths and sequencing data amounts on the calculation of transcript expression. EdgeR software (Robinson et al., 2010) was employed to screen differentially expressed lncRNAs (DElncRNAs) in the AmCK1 vs AmT1, AmCK2 vs AmT2, and AmCK3 vs AmT3 comparison groups following the criteria of *P*≤0.05 (corrected by false discovery rate) and |log_2_(FC)|≥1. Finally, Venn diagram analysis and expression clustering of DElncRNAs in different comparison groups were conducted using the related tools in the OmicShare platform (www.omicshare.com).

### Investigation of the *cis*-acting role of DElncRNAs

Protein-coding genes located 10 kb upstream and downstream of DElncRNAs were surveyed and then annotated to GO (http://www.geneontology.org/) and KEGG (https://www.kegg.jp/) databases utilizing the Blast tool. Subsequently, gene numbers were calculated for each GO term or KEGG pathway. Furthermore, significantly enriched GO terms by neighboring genes were defined by hypergeometric testing, while KEGG pathway enrichment analysis was performed by KOBAS 2.0 software (Xie et al., 2011), with the *A. mellifera* reference genome (Amel_HAV3.1) as the background. Only terms or pathways with corrected *p* values of less than 0.05 were considered enriched.

### Source of small RNA-seq datasets

In our previous study, gut tissues of *A. apis*-inoculated 4-, 5-, and 6-day-old larvae and corresponding uninoculated 4-, 5-, and 6-day-old larval guts were prepared following the above mentioned method, followed by total RNA isolation, cDNA library construction, and sRNA-seq by Guangzhou Gene Denovo Biotechnology Co., Ltd. using the Illumina MiSeq™ 4000 platform. Raw data derived from sRNA-seq are available in the NCBI SRA database under the BioProject number PRJNA406998. Quality control was previously performed, and the results were suggestive of the high quality of sRNA-seq-derived data (Fen et al., 2022).

### Investigation of ceRNA regulatory networks

According to the method described by Chen et al. (2019a), target DEmiRNAs of DElncRNAs and target DEmRNAs of DEmiRNAs were predicted by using a combination of three software programs, MiRanda (V3.3a), RNAhybrid (V2.1.2) + SVM_light (V6.01), and TargetFind, and the intersection was deemed reliable targets. Based on the prediction results, DElncRNA-DEmiRNA-DEmRNA regulatory networks were constructed and visualized by Cytoscape software (Smoot et al., 2011) with default parameters.

### RT-qPCR validation of DElnRNAs

Nine DElncRNAs were randomly selected for RT-qPCR, including three (MSTRG.1133.1, XR_001704875.2, and MSTRG.11613.1) from the AmCK1 vs AmT1 comparison group, three (MSTRG.4918.2, MSTRG. 9603.5, and MSTRG.8790.2) from AmCK2 vs AmT2 comparison group and three (XR_001705688.2, XR_410074.3, and XR_003304187.1) from AmCK3 vs AmT3 comparison group. Specific forward and reverse primers for each of the nine selected DElncRNAs were respectively designed using Primer Premier 6, and synthesized by Sangon Biotech (Shanghai) Co., Ltd. Gene *actin* (GeneBank ID: 406122) was used as internal reference. Total RNA from gut samples in each group were respectively isolated using FastPure Cell/Tissue Total RNA Isolation Kit V2 (Vazyme, China), followed by reverse transcription with Random primers. The resulting cDNA were used as templates for qPCR reaction, which was performed on QuanStudio 3 Fluorescence quantitative PCR instrument (ABI, USA) The reaction system (20 μL) contained 10 μL of SYBR Green Dye, 1 μL of upstream and downstream primers(10 μmol/L), 1 μL of cDNA template, 7 μL of DEPC water. The reaction conditions were set as follows: 95°C pre-denaturation for 5 min; 95°C denaturation for 30 s, 60°C annealing and extension for 30 s, a total of 40 cycles each group of qPCR reaction and set three times experiment for repeating. The relative expression level of each DElncRNA was calculated using 2^-ΔCt^ method (Livak et al., 2001). All experiments were run with at least three parallel samples and were repeated three times. Data were shown as mean ± standard deviation (SD) and subjected to Student’s *t*-test by Graph Prism 8 software (*P*<0.05 was considered statistically significant). Detailed information of primers used in this work was presented in the Table S1.

## Result

### Quality control of deep sequencing data

Totally, 85 811 046, 81 962 296, 85 636 572, 79 267 686, 82 889 882, and 100 211 796 raw reads were produced from AmCK1, AmCK2, AmCK3, AmT1, AmT2, and AmT3 groups, respectively (Table S2). After strict quality control, 85 739 414, 81 896 402, 85 573 798, 79 202 304, 82 828 926, and 100 128 692 clean reads were obtained, respectively (Table S2). In brief, the Q20 and Q30 of clean reads in six groups were above 98.07% and 94.30%, respectively (Table S2). In addition, the mapping ratio of clean reads in the reference genome was above 99.92% (Table S2). The results showed that the next-generation sequencing data were reliable and could be used for downstream analyses. Compared with the *A. m. ligustica* reference genome (Amel_HAV3.1), the comparison rate ranged from 92.06% to 94.81%, and there were 63.10% - 69.80% clean reads were compared to the exon region, 8.56% - 9.59% of them compared to the intron region, and 20.98% - 27.32% compared to the intergene region (Table S3).

### Identification and investigation of *A. m. ligustica* lncRNAs

In the aforementioned six groups, 1 991, 2 031, 1 970, 2 101, 1 955, and 1 944 lncRNAs were identified. After removing redundant lncRNAs, a total of 3 746 *A. m. ligustica* lncRNAs were discovered, including 3 146 known lncRNAs and 600 novel lncRNAs. Among these, there were 78 sense lncRNAs, 891 antisense lncRNAs, 1 893 intergenic lncRNAs, 346 bidirectional lncRNAs, and 210 intron lncRNAs (Figure 1).

**Figure 1.**
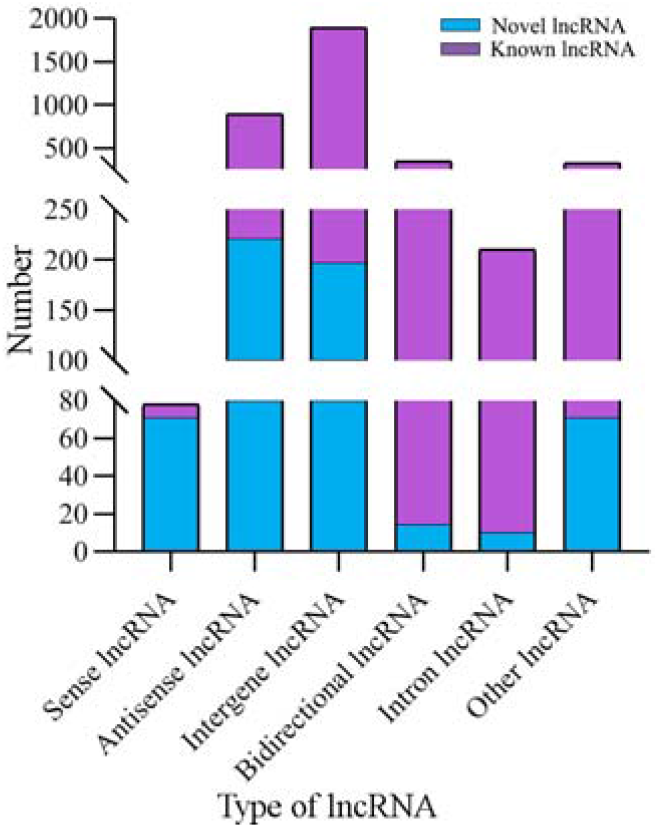
Type statistics of *A. m. ligustica* lncRNAs identified in this work

### Differential expression profile of lncRNAs in larval guts infected by *A. apis*

In the AmCK1 vs AmT1 comparison group, 156 upregulated and 201 downregulated lncRNAs were identified; 98 upregulated and 138 downregulated lncRNAs were identified in the AmCK2 vs AmT2 comparison group; and 361 upregulated and 144 downregulated lncRNAs were identified in the AmCK3 vs AmT3 comparison group (Figure 2 A). Additionally, Venn analysis indicated that 32 DElncRNAs were shared by the three comparison groups mentioned above, and the numbers of specific DElncRNAs were 191, 93, and 322, respectively (Figure 2 B). As shown in Figure 2 C, the shared DElncRNAs displayed various expression patterns during *A. apis* infection; such as the expression levels of XR_003305757.1, XR_003304308.1, and XR_001705213.2 were elevated in AmCK1 group but decreased in AmT1 group(Figure 2 C).

**Figure 2.**
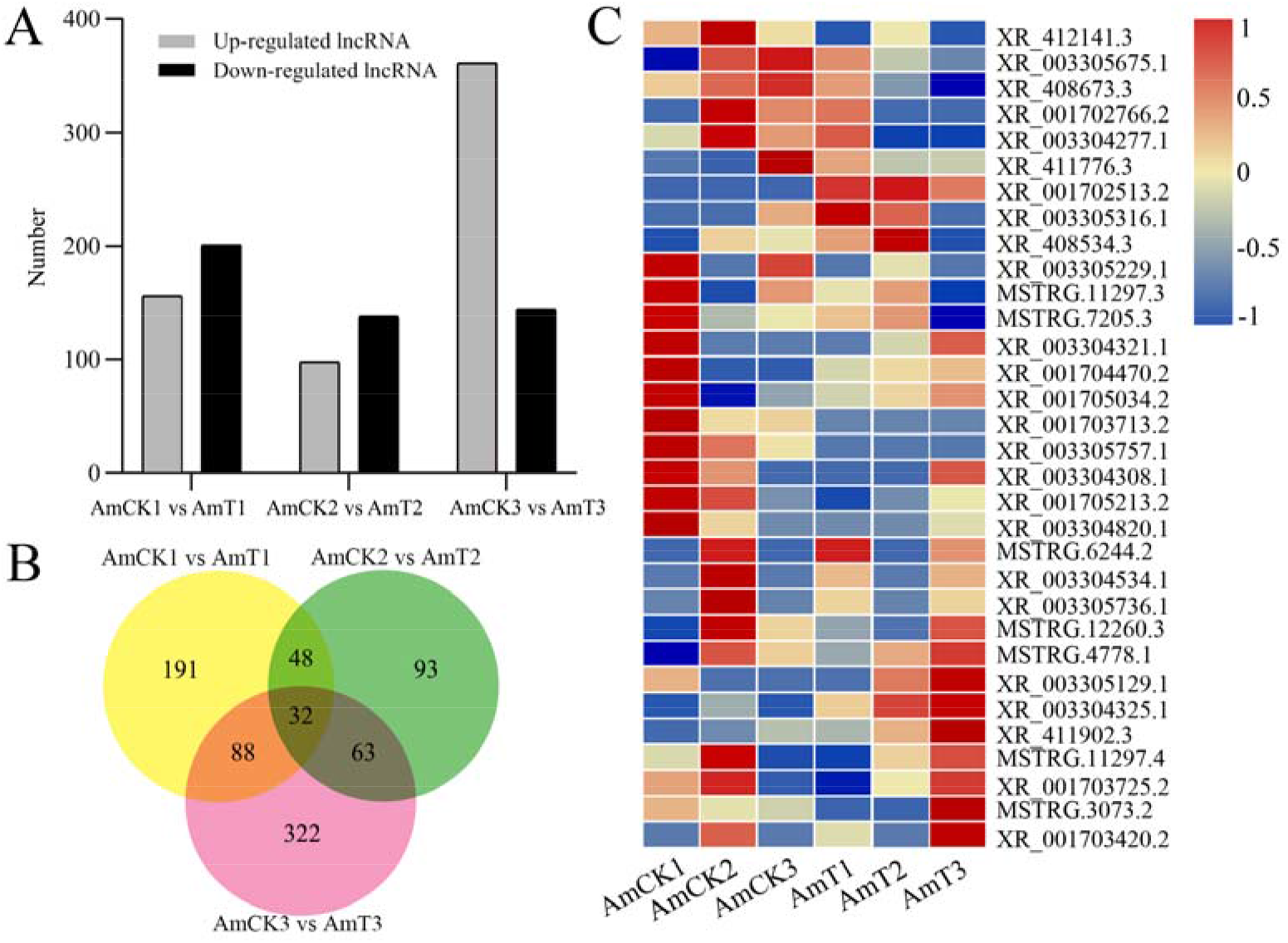
Number and expression of DElncRNAs in *A. m. ligustica* larval guts responding to *A. apis* infection A: Number of up- and down-regulated lncRNAs in three comparison groups; B: Venn diagram of DElncRNAs in three comparison groups; C: Heat map of shared DElncRNAs by three comparison groups

**Figure 2.**
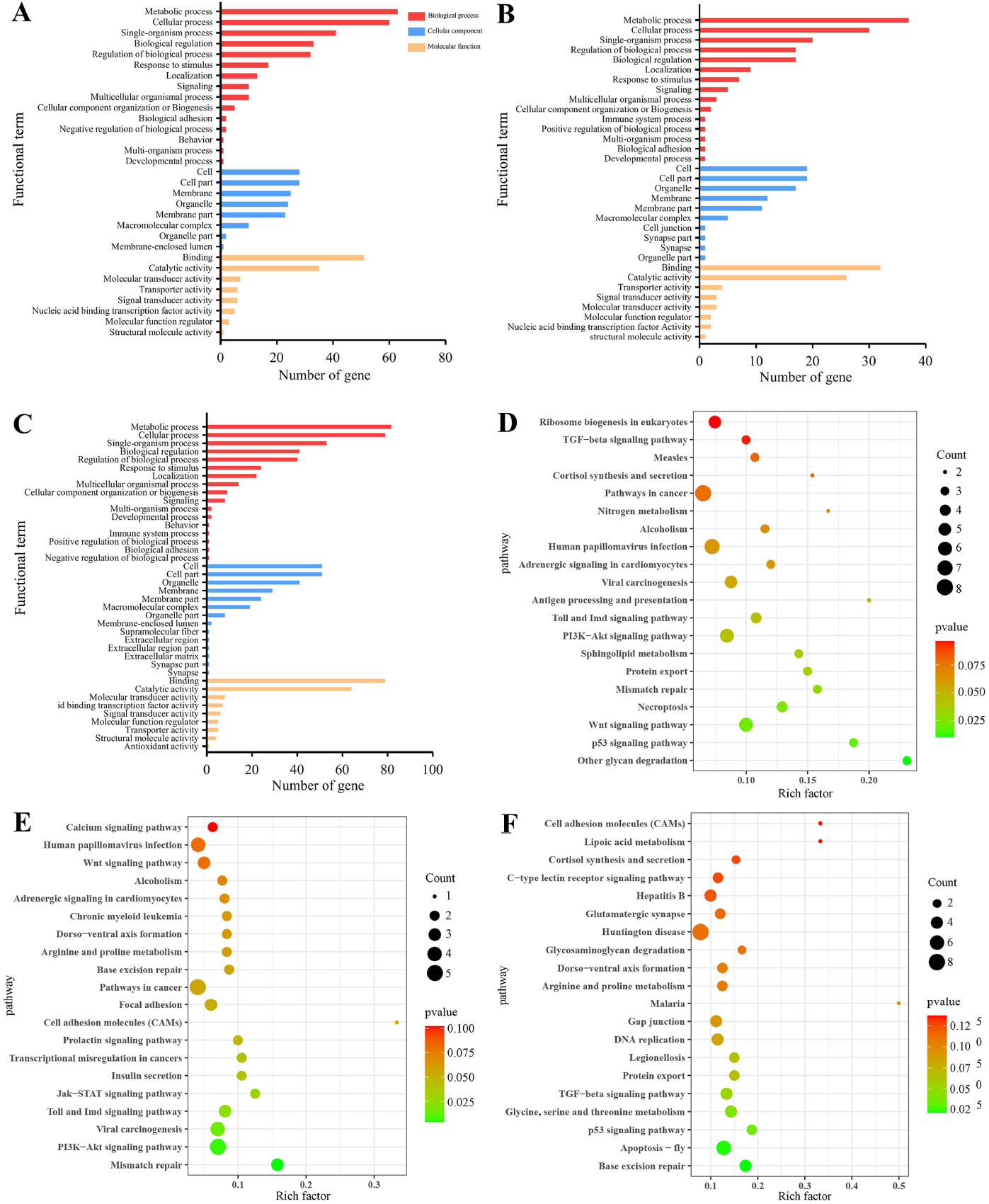
Number and expression of DElncRNAs in *A. m. ligustica* larval guts responding to *A. apis* infection

### C*is*-acting regulation of host DElncRNAs

In the AmCK1 vs AmT1 comparison group, 217 DElncRNAs were predicted to regulate 361 neighboring genes, which were enriched in 31 GO terms relative to biological process, cellular component, and molecular function, such as behavior, cell, and binding (Figure 3 A); these genes were also involved in 203 KEGG pathways, including ribosome biogenesis in eukaryotes, measles, and viral carcinogenesis (Figure 3 D); further investigation suggested that 37 genes were associated with immune pathways such as cell apoptosis, autophagy, and endocytosis (Table 1). Comparatively, 129 DElncRNAs in the AmCK2 vs AmT2 comparison group were predicted to regulate 217 neighboring genes, which were enriched in 33 GO terms (Figure 3 B), such as localization and organelle, as well as 154 KEGG pathways, such as alcoholism and cell adhesion molecules (Figure 3 E); additionally, 19 genes relevant to immune pathways, such as phagosome, ubiquitin-mediated proteolysis, and Jak-STAT signaling pathway, were putatively regulated by 129 DElncRNAs (Table 1). In the AmCK3 vs AmT3 comparison group, 272 DElncRNAs were predicted to regulate 496 neighboring genes, which were enriched in 31 GO terms (Figure 3 C), such as immune system process and catalytic activity, as well as 226 KEGG pathways, such as malaria and p53 signaling pathway (Figure 3 F); 272 DElncRNAs were found to be engaged in regulating 51 genes related to cellular or humoral immune pathways, such as endocytosis, biosynthesis of insect hormones, and lysosome (Table 1).

**Figure 3.**
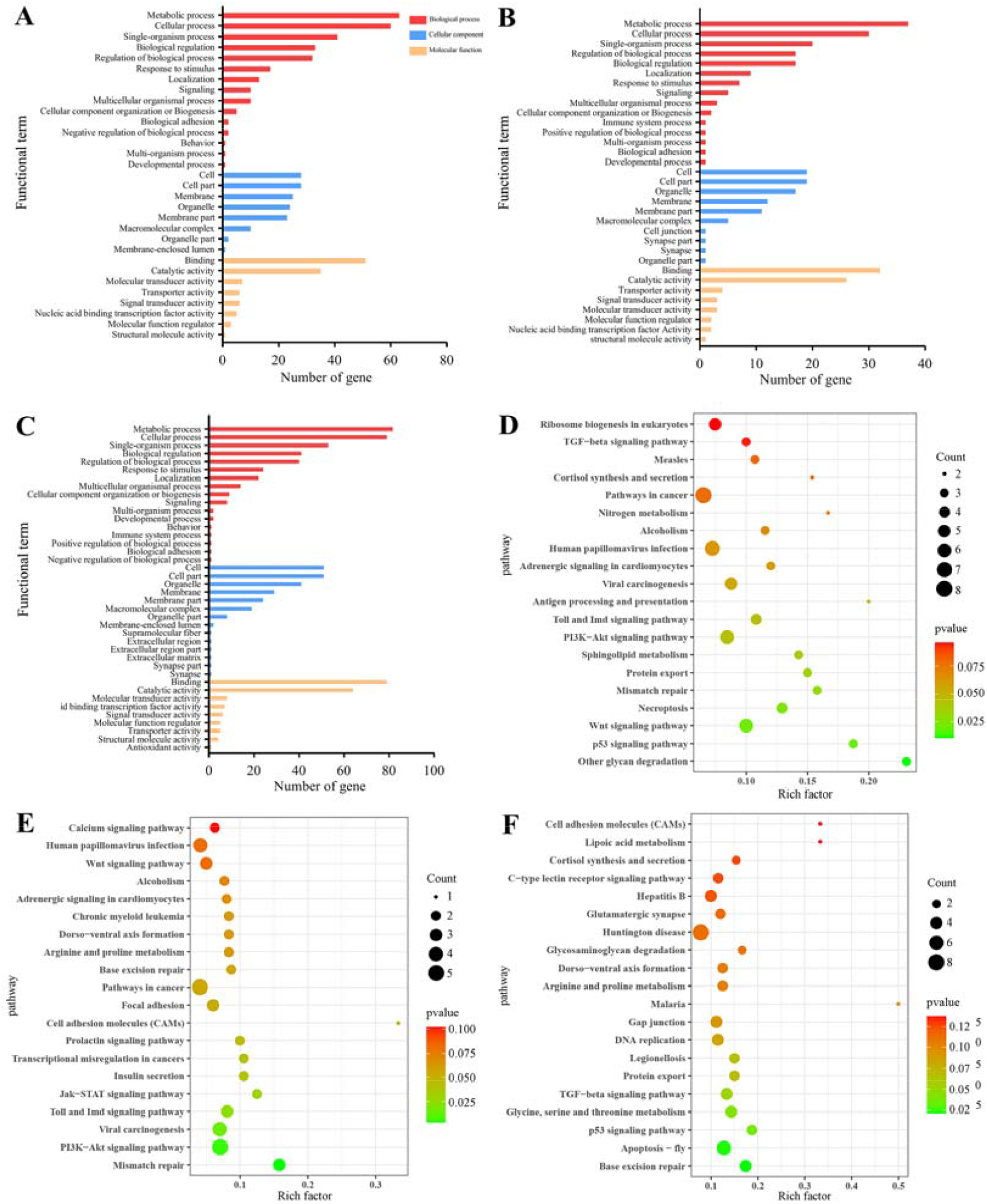
GO terms and KEGG pathways enrich by neighboring genes of DElncRNAs A-C: GO classification of neighboring genes of DElncRNAs in AmCK1 vs AmT1, AmCK2 vs AmT2, and AmCK3 vs AmT3 comparison groups; E-F: Enriched KEGG pathways by neighboring genes of DElncRNAs in AmCK1 vs AmT1, AmCK2 vs AmT2, and AmCK3 vs AmT3 comparison groups

**Table 1.** Summary of neighboring genes of DElncRNAs associated with immune pathways

|  |  |  |  |
| --- | --- | --- | --- |
| Pathway | AmCK1 vs AmT1<br>comparison group | AmCK2 vs AmT2<br>comparison group | AmCK3 vs AmT3<br>comparison group |
| Cell apoptosis | 3 | 1 | 10 |
| Cell apoptosis-multi-species | 1 | 0 | 2 |
| Autophagy-animal | 1 | 0 | 3 |
| Endocytosis | 3 | 1 | 6 |
| Biosynthesis of insect hormones | 0 | 0 | 1 |
| Lysosome | 3 | 2 | 4 |
| Necroptosis | 4 | 0 | 2 |
| Phagosome | 2 | 2 | 4 |
| Ubiquitin-mediated proteolysis | 3 | 2 | 4 |
| Jak-STAT signaling pathway | 2 | 2 | 1 |
| MAPK signaling pathway | 7 | 4 | 7 |
| NF-κB signaling pathway | 1 | 0 | 0 |
| Ras signaling pathway | 2 | 2 | 3 |
| Toll and Imd signaling pathway | 4 | 3 | 3 |
| Toll-like receptor signaling pathway | 1 | 0 | 1 |

### CeRNA regulatory network of host DElncRNAs

Complex ceRNA networks of DElncRNAs existed in the AmCK1 vs AmT1 and AmCK2 vs AmT2 comparison groups, whereas the DElncRNA-DEmiRNA-DEmRNA regulatory network in the AmCK3 vs AmT3 comparison groups was more complicated (Figure 4). In detail, 197 DElncRNAs in the AmCK1 vs AmT1 comparison group could target 10 DEmiRNAs and further bind to 147 DEmRNAs (Figure 4 A), which were engaged in 24 GO terms, including single-organism process and membrane (Figure 5 A), as well as 23 KEGG pathways, such as circadian rhythm-fly and dorsoventral axis formation (Figure 5 D). In the AmCK2 vs AmT2 comparison group, 95 DElncRNAs could target eight DEmiRNAs and further link to 79 DEmRNAs (Figure 4 B), which were involved in 22 terms, including single-organism process and binding (Figure 5 B), as well as 16 pathways, such as ECM-receptor interaction and beta-alanine metabolism (Figure 5 E). In the AmCK3 vs AmT3 comparison group, 356 DElncRNAs could target 21 DEmiRNAs and 315 DEmRNAs (Figure 4 C), which were associated with 28 terms, including cellular process and membrane part (Figure 5 C), as well as 68 pathways, such as neuroactive ligand □ receptor interaction and drug metabolism-other enzymes (Figure 5 F).

**Figure 4.**
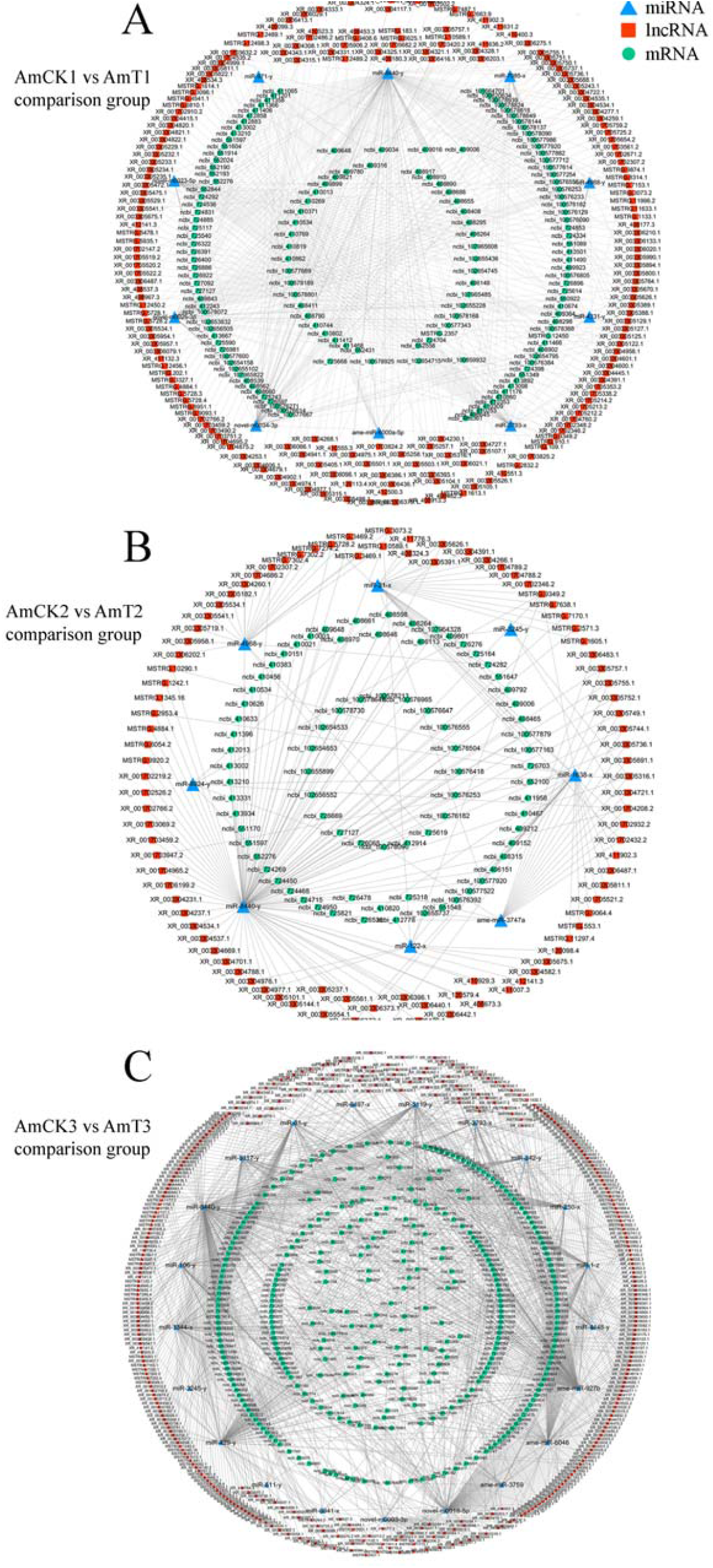
DElncRNA-DEmiRNA-DEmRNA regulatory networks involved in response of *A. m. ligustica* larval guts to *A. apis* infection

**Figure 5.**
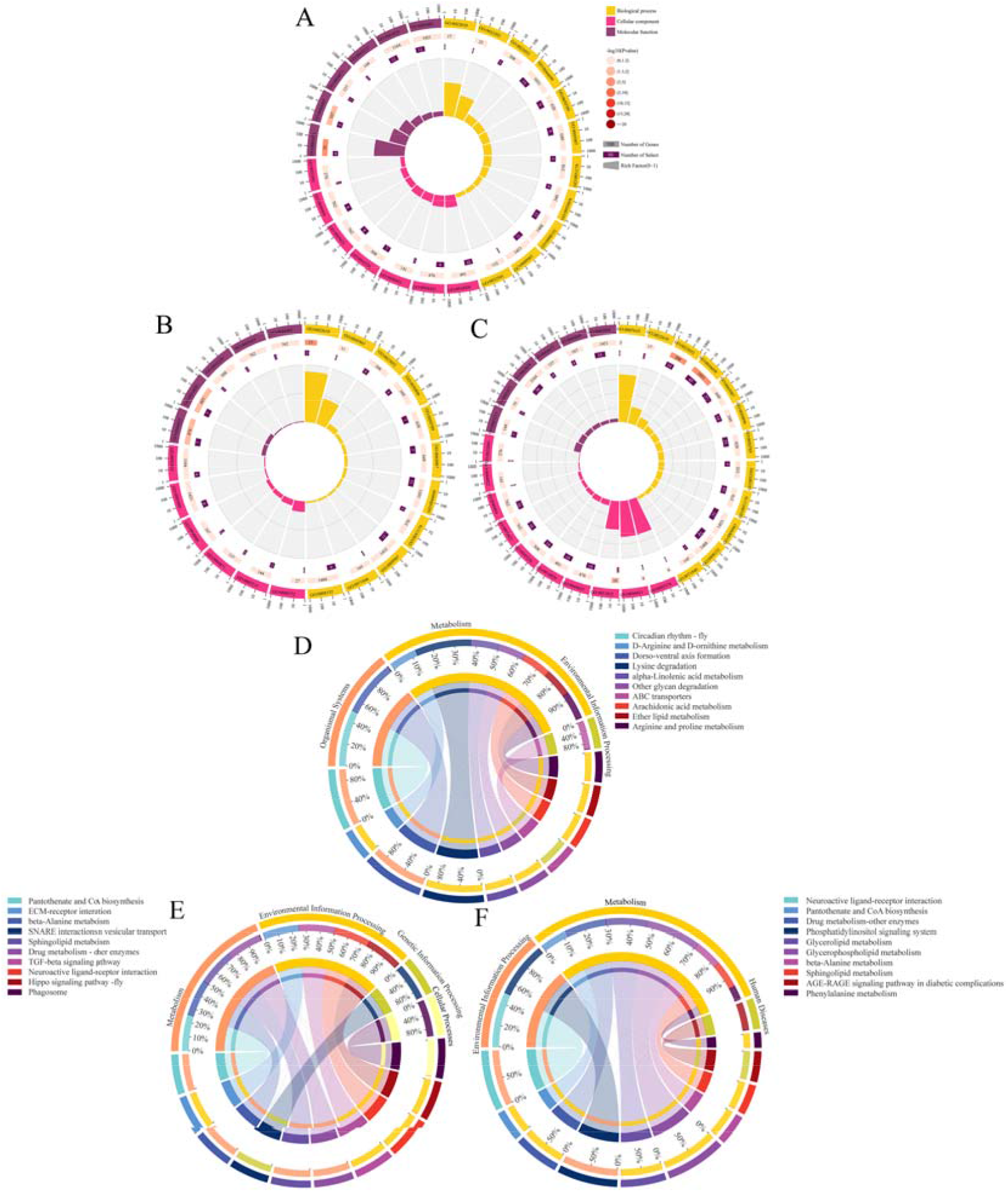
GO terms and KEGG pathways enriched by target DEmRNAs within ceRNA networks in three comparison groups A-C: Loop graphs of enriched GO terms by targets in AmCK1 vs AmT1, AmCK2 vs AmT2, and AmCK3 vs AmT3 comparison groups; D-F: Chord graphs of enriched KEGG pathways by targets in AmCK1 vs AmT1, AmCK2 vs AmT2, and AmCK3 vs AmT3 comparison groups

Further investigation demonstrated that target DEmRNAs in the AmCK1 vs AmT1 comparison group were related to two cellular immune pathways (lysosome and endocytosis) and one humoral immune pathway (MAPK signalinging pathway); target DEmRNAs in the AmCK2 vs AmT2 comparison group were associated with two cellular immune pathways (endocytosis and phagosome); target DEmRNAs in the AmCK3 vs AmT3 comparison group were relevant to six cellular immune pathways (lysosome, endocytosis, phagosome, etc.) and one humoral immune pathway (Toll and Imd signalinging pathways). Detailed information about immune-related targets is presented in Table 2.

**Table 2.** Summary of immune-associated target DEmRNAs within ceRNA networks

| Pathways | AmCK1 vs AmT1<br>comparison group | AmCK2 vs AmT2<br>comparison group | AmCK3 vs AmT3<br>comparison group |
| --- | --- | --- | --- |
| Lysosome | 1 | 0 | 1 |
| Endocytosis | 1 | 1 | 2 |
| Phagosome | 0 | 1 | 1 |
| Autophagy-animals | 0 | 0 | 4 |
| Apoptosis-fly | 0 | 0 | 2 |
| Ubiquitin-mediated proteolysis | 0 | 0 | 2 |
| MAPK signaling pathway-fly | 1 | 0 | 1 |
| Toll and Imd signaling pathway | 0 | 0 | 2 |

### Validation of DElncRNAs by RT□qPCR

The RT□PCR results suggested that the expression trends of nine DElncRNAs were consistent with those in the transcriptome data (Figure 6), further confirming the reliability of the sequencing data used in this work.

**Figure 6.**
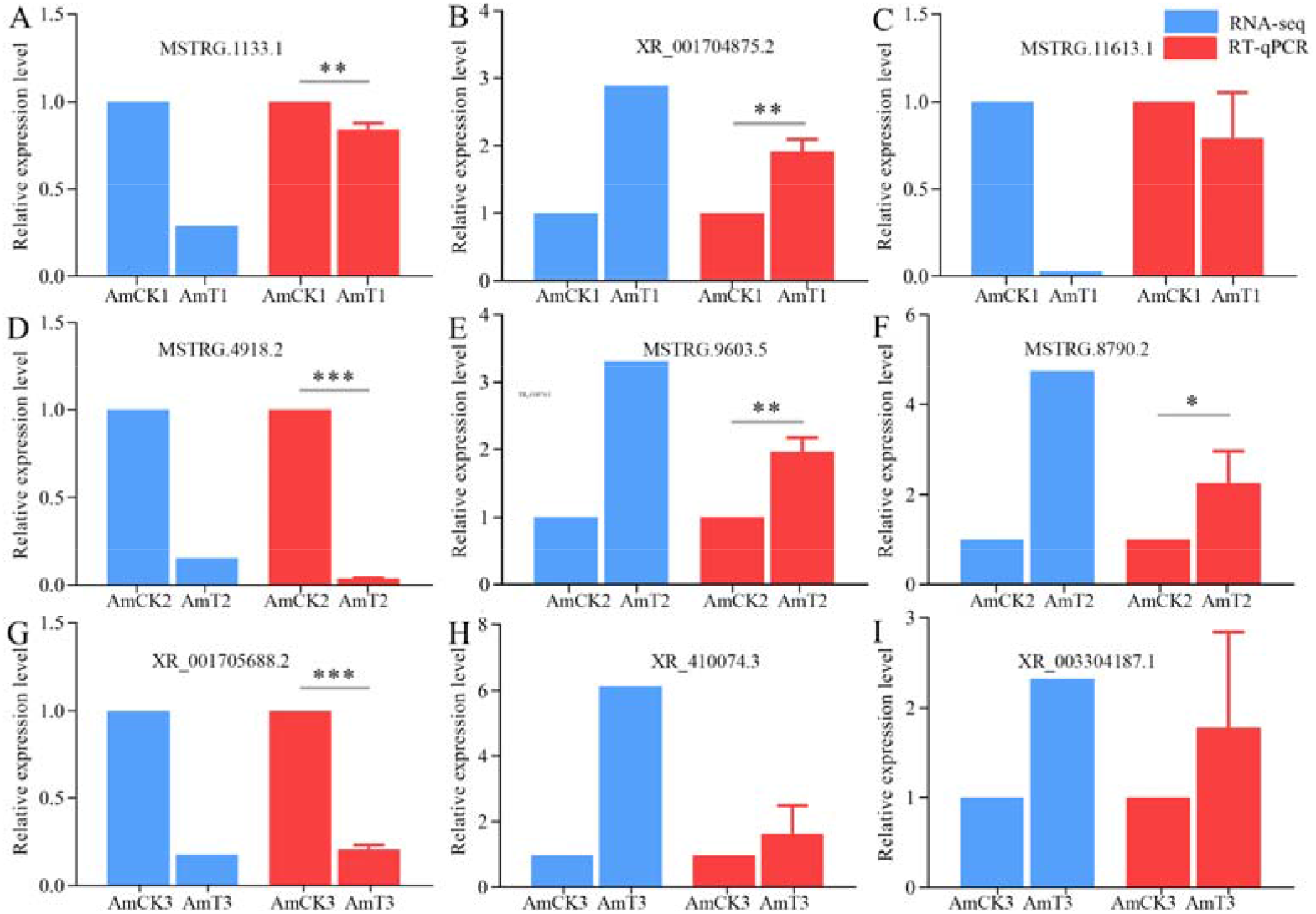
RT-qPCR verification of DElncRNAs * represents *P*<0.05, ** represents *P*<0.01, *** represents *P*<0.001

**Figure 7.**
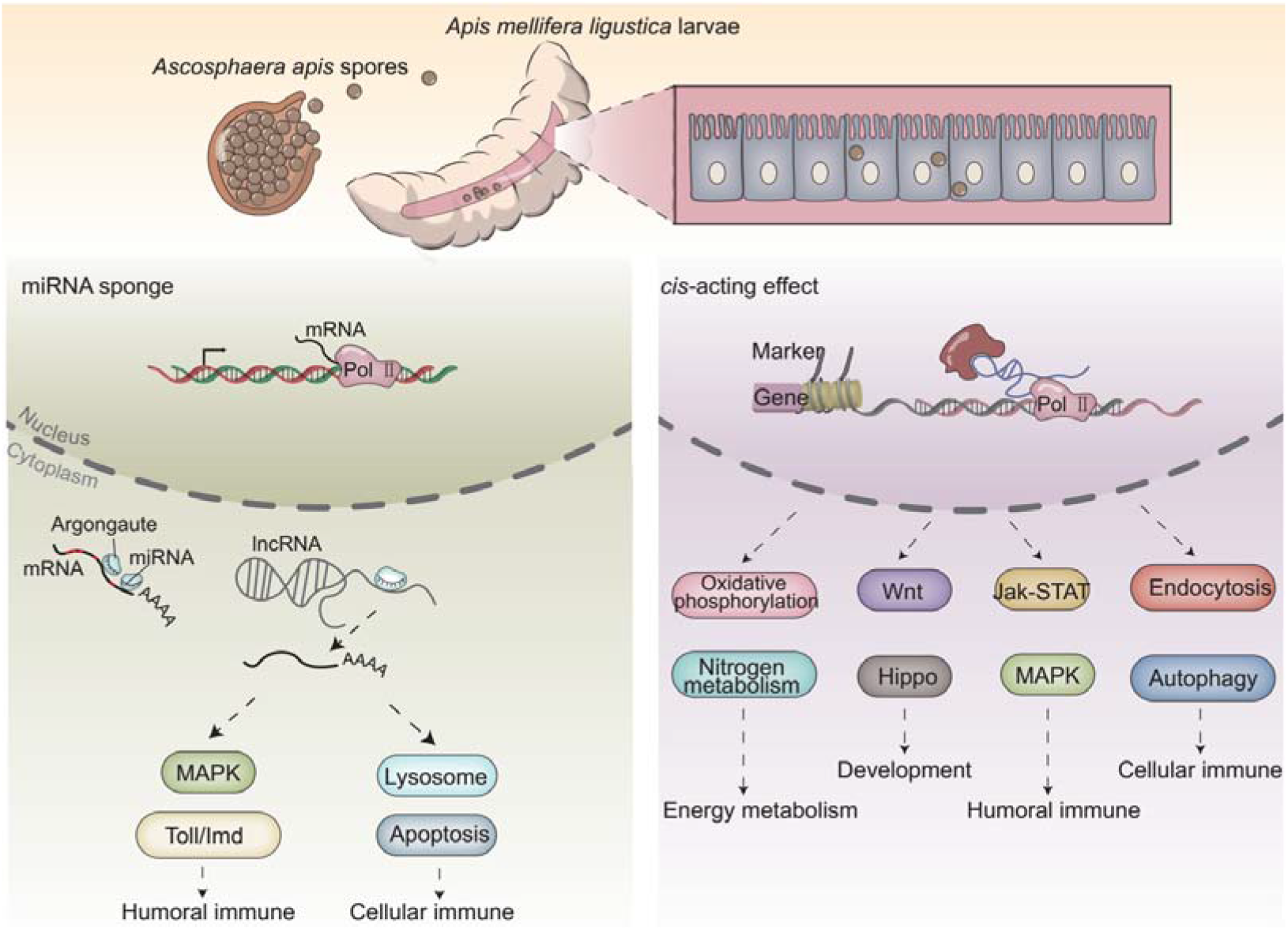
A working model of DElncRNA-modulated larval immune response of *A. m. ligustica* bee to *A. apis* invasion

## Discussion

Previously, our group identified 6 353 lncRNAs in *A. m. ligustica* workers’ midguts based on RNA-seq, including 4 749 known and 1 604 novel lncRNAs (Chen et al., 2019a). In the current work, a comprehensive investigation and the potential regulatory function of lncRNAs in the larval response of *A. m. ligustica* to *A. apis* invasion were conducted for the first time, and 3 146 known and 600 novel lncRNAs were identified (Figure 1). Given the limited number of honey bee-derived lncRNAs, the identified lncRNAs could further enrich the reservoir of honey bee lncRNAs and offer valuable genetic resources for further functional study. We compared lncRNAs discovered in this work with those identified in workers’ midguts and found that 39 (1.2%) known lncRNAs and no novel lncRNAs were shared, indicating that the shared known lncRNAs may play fundamental roles in both the adult bee midgut and larval gut, whereas those specific lncRNAs may have different functions at different developmental stages of the gut tissue. In animals and plants, lncRNA expression is tissue- and stage-specific (Statello et al., 2021). Hence, the total number of *A. m. ligustica* lncRNAs should be much greater than the documented lncRNAs of this research.

Here, 357, 236, and 505 lncRNAs were differentially expressed in 4-, 5-, and 6-day-old larval guts inoculated with *A. apis* compared with uninoculated larval guts (Figure 2 A), suggesting that a portion of lncRNAs were induced to activation while some other ones were suppressed by *A. apis*. These DElncRNAs were speculated to be engaged in the host response. This was also indicative of the overall alteration of host lncRNAs occurring due to *A. apis* infection. In addition, three up-regulated lncRNAs, XR_001702513.2 (log_2_FC=8.748 2, *P*=0.000 2), XR_003304325.1 (log_2_FC=1.854 1, *P*=0.000 4), and XR_411902.3 (log_2_FC=2.667 4, *P*<0.000 1) and two down-regulated lncRNAs, XR_001703713.2 (log_2_FC=-7.965 8, *P*<0.000 1) and XR_003305757.1 (log_2_FC=-7.409 4, *P*<0.000 1) were shared by the above mentioned three comparison groups. (Figure 2 C). It is inferred that these five shared DElncRNAs played critical roles in the *A. apis* response of larval guts, deserving additional functional investigation.

Accumulating evidence has shown that lncRNAs are capable of regulating the transcription of neighboring genes in a *cis*-acting manner (Gil and Ulitsky, 2020). Chen et al. (2019a) constructed DElncRNA-miRNA-mRNA networks of the *A. m. ligustica* response to *N. ceranae* infection and found that a portion of DElncRNAs were likely to participate in regulating host material and energy metabolism as well as cellular and humoral immunity during host responses to *N. ceranae* invasion. Ropri et al. (2021) reported that SE-lncRNAs (RP11-379F4.4 and RP11-465B22.8) played a potential role in the progression of ductal carcinoma in situ (DCIS) and invasive ductal carcinoma (IDC) by regulating the expression of neighboring genes. Insects only consume a small amount of energy to maintain basic activities without activation of the immune system; however, once the immune system is activated under pathogen invasion, the energy consumption rate will greatly increase (Kingsolver et al., 2013). Here, it was observed that the largest number of neighboring genes in the AmCK3 vs AmT3 comparison group were involved in energy metabolism-associated pathways such as oxidative phosphorylation and nitrogen metabolism, suggestive of the continuous proliferation of *A. apis* within the host larval gut and the *A. apis*-caused pressure at the later stage of infection, which resulted in the enhancement of host immune defense and thus the elevation of the energy metabolism rate. To cope with the infection of pathogenic microorganisms, insects have evolved an efficient innate immune system, which can be divided into cellular immune and humoral immune systems. The former mainly includes phagocytosis, nodules, encapsulation, and melanization of hemolymph, while the latter is mediated by multiple signalinging pathways, such as the Toll, Imd, and JAK/STAT signalinging pathways (Hillyer, 2016; Valanne et al., 2011). The Toll signalinging pathway plays a major role in the immune defense and development of insects. The Imd signalinging pathway regulates the expression of genes encoding antimicrobial peptides and the synthesis of proteins (Myllymäki et al., 2014). Here, 16, eight, and 29 neighboring genes of DElncRNAs in the guts of *A. apis*-inoculated 4-, 5-, and 6-day-old larvae were associated with a subseries of cellular immune pathways, such as cell apoptosis, cell apoptosis, and autophagy; additionally, 10, six, and eight targets were related to several humoral immune pathways, such as the Jak-STAT, MAPK, and NF-κB signalinging pathways (Figure 3). These results demonstrated that corresponding DElncRNAs were likely to modulate the transcription of neighboring genes and further participate in regulating host cellular and humoral immune responses to *A. apis* infection.

Those lncRNAs with MREs could interact with miRNAs and influence downstream gene expression via ceRNA networks (Tang et al., 2017; Wang et al., 2020). An increasing number of lncRNAs have been verified to be pivotal regulators in the occurrence and development of an array of human diseases through ceRNA mechanisms, such as cancer, Alzheimer’s disease, and cardiovascular disease (Wu et al., 2020; Ma et al., 2021). Previous findings showed that lncRNA-mediated ceRNA regulatory networks were putatively engaged in midgut development and the *Nosema ceranae* response of western honey bee workers (Guo et al., 2018; Chen et al., 2019a). In this work, 197, 95, and 356 DElncRNAs in 4-, 5-, and 6-day-old larval guts could respectively target 10, eight, and 21 DEmiRNAs and further target 147, 79, and 315 DEmRNAs, forming complex ceRNA regulatory networks (Figure 4). Additionally, four, four, and 14 DEmRNAs within ceRNA networks were relevant to an immune-related GO term, response to the stimulus; two, two, and 12 DEmRNAs could be annotated to six cellular immune pathways, namely, lysosome, endocytosis, and apoptosis. Together, these results demonstrated that corresponding DElncRNAs (ncbi_410614, ncbi_412360, and ncbi_409522, etc.) were putative regulators in the aforementioned cellular immune pathways and response to the stimulus. Intriguingly, three DEmRNAs in the AmCK3 vs AmT3 comparison group were involved in two humoral immune pathways, including the MAPK and Toll and Imd signalinging pathways. This indicated that a portion of DElncRNAs (ncbi_725117, ncbi_406086, ncbi_100302584, etc.) in host guts potentially exerted regulatory functions at the later stage of *A. apis* invasion.

miR-1 plays an important role in the pathogenesis of heart disease, Liu et al. (2021) indicated that inhibiting miR-1 may relieve right ventricle hypertrophy and fibrosis in model rats used in their research, which also works significantly in plants. Wang et al. (2018) discovered that Connexin43, an extract of *Astragalus* root, worked by targeting miR-1 to cure viral myocarditis, and overexpression of miR-1 inhibited endogenous Connexin43 expression significantly. Previous studies have found that the expression of miR-1 in *A. m. ligustica* workers was significantly downregulated 6 days after *N. ceranae* inoculation, suggesting its potential involvement in the host immune response (Huang et al., 2015). In the Asian honey bee *A. ceranae*, Chen et al. (2019b) found that the expression of miR-1-x in the worker’s midgut was significantly downregulated at 7 d post inoculation with *N. ceranae* spores. Here, 35 DElncRNAs in the 6-day-old larval gut could jointly target miR-1-z (highly homologous to ame-miR-1), which can further target 32 DEmRNAs (Figure 4 C). It is speculated that these DElncRNAs potentially regulate the expression of downstream target genes by targeting miR-1-z, further modulating the larval immune response to *A. apis* invasion. Effective knockdown of lncRNAs in insects such as *Drosophila, Helicoverpa armigera*, and *Plutella xylostella* was achieved utilizing the RNAi method (Zhang et al., 2020; Guan et al., 2020). Recently, our team knocked down lncRNA13164 in *A. ceranae* larval guts based on dsRNA-based dsRNA and revealed that lncRNA13164 regulated the expression of three immune genes (*stk, e3μl* and *or1*) via ace-miR-4968 and further mediated the host immune response to *A. apis* invasion (Fu et al., 2022). In the near future, we will perform a functional study on miR-1-z as well as associated DElncRNAs and explore the mechanism underlying the DElncRNA-miR-1-z-DEmRNA axis-regulated host response.

## Conclusions

In a nutshell, 3 146 known lncRNAs and 600 novel lncRNAs were identified in the *A. mellifera* larval guts; additionally, *A. apis* infection caused overall change of expression profile of lncRNAs in host guts; DElncRNAs potentially participated in larval immune response to *A. apis* invasion by regulating the expression of neighboring genes or interacting with DEmiRNAs; corresponding DElncRNAs were potentially engaged in host immune response through ceRNA regulatory networks via absorption of miR-1-z.

## Data availability statement

The raw data supporting the conclusion of this article will be made available by the authors, without undue reservation.

## Ethics statement

The study was reviewed and approved by Fujian Agriculture and Forestry University.

## Author contributions

RG, XF, and JW conceived and planned the experiments. YY, QL, WZ, and ZC carried out the experiments, analyzed and interpreted the data. RG, MS, XG, and PZ designed the figures. DC and RG reviewed and edited the paper. All authors contributed to the review and approval of the manuscript for publication.

## Acknowledgments

All authors thanks reviewers and editors for their constructive comments and recommendations. RG appreciates the love from his beloved wife and daughter.

## Publisher’s note

All claims expressed in this article are solely those of the authors and do not necessarily represent those of their affiliated organizations, or those of the publisher, the editors and the reviewers. Any product that may be evaluated in this article, or claim that may be made by its manufacturer, is not guaranteed or endorsed by the publisher.

## Supplementary material

The Supplementary Material for this article can be found online at: https://www.frontiersin.org/ Table S1, Table S2, and Table S3.

